# Prediction transfer does not imply reliability transfer in single-cell foundation models

**DOI:** 10.64898/2026.10.21.760716

**Authors:** Aditya Sriram, Tianze Luo, Laizhi Zhang, Soyeon Kim, Hyun Jung Park

## Abstract

Single-cell foundation models are increasingly transferred across tissues, diseases, perturbation screens, and experimental platforms. Although substantial effort has focused on prediction transfer, it remains unclear whether a model’s assessment of its own reliability transfers similarly across biological contexts. Here we define reliability transfer as the ability to transfer an accurate assessment of when model predictions are likely to be correct or incorrect and investigate its behavior across ten perturbation-prediction systems spanning multiple datasets, models, and evaluation settings. We find that reliability transfer varies substantially across source-target pairs and cannot be inferred from predictive performance alone, indicating that prediction transfer and reliability transfer constitute distinct forms of generalization. Analyses of biological annotations, network connectivity, target exposure, and contextual factors reveal that prediction reliability is associated with both shared and context-dependent signals, with their relative importance differing across systems. To characterize this variability, we introduce reliability-attribution fingerprints, context-level signatures derived from strictly cross-fitted SHAP explanations that summarize how reliability estimators combine biological and contextual information when assigning confidence. Across systems, contexts with similar attribution-magnitude fingerprints tended to transfer reliability more successfully, and fingerprint similarity identified transfer sources achieving higher reliability-transfer performance within the observed candidate set. However, these relationships weakened under stricter cross-family transfer settings, indicating that transferable reliability structure remains partly dependent on shared contextual information. Finally, transferred reliability estimators produced different perturbation rankings despite identical perturbation predictions, illustrating a practical consequence of reliability transfer for downstream prioritization. Together, these results establish reliability transfer as a distinct aspect of foundation-model generalization and introduce reliability-attribution fingerprints as a framework for characterizing transferable reliability structure across biological contexts.

## Introduction

As single-cell foundation models increasingly shape biological discovery, their value depends not only on transferring predictive knowledge to new contexts but also on determining when those predictions can be trusted. Existing efforts have therefore focused primarily on prediction transfer, namely the ability of a model trained in one biological context to maintain predictive performance in another ^1, 2^. Foundation models are routinely transferred across tissues, diseases, perturbation screens, and experimental platforms, enabling knowledge learned from large reference datasets to be reused in settings where labeled data are limited ^2–4^. However, successful prediction transfer does not necessarily imply successful reliability transfer, the ability to transfer an accurate assessment of when model predictions are likely to be correct or incorrect. Just as predictive accuracy and calibration capture distinct aspects of model performance within a dataset ^5, 6^, prediction transfer and reliability transfer capture distinct aspects of model generalization across datasets. Consequently, two transferred reliability estimators can assign different levels of confidence to the same predictions, potentially leading to different perturbations being prioritized for downstream evaluation despite comparable predictive performance. In such cases, prediction transfer remains successful, yet transferred reliability estimates can support different prioritization decisions. As foundation models, reference datasets, and transfer-learning strategies continue to proliferate ^2, 3^, understanding when reliability can be transferred across biological contexts has become an important challenge for trustworthy and efficient AI-guided biological discovery.

Existing approaches provide powerful tools for addressing individual aspects of this challenge. Model calibration ^6, 7^, uncertainty estimation ^8^, conformal prediction ^9^, and attribution-based methods ^10, 11^ can assess different aspects of model confidence, uncertainty, coverage, or feature contributions within a dataset, while transfer-learning frameworks have demonstrated how predictive knowledge can be reused across tissues, diseases, and perturbation settings ^1, 2, 4^. Specifically, calibration examines whether confidence agrees with observed frequencies of correctness, uncertainty estimation characterizes predictive uncertainty, conformal prediction constructs prediction sets or intervals with coverage guarantees under stated assumptions, and attribution methods explain how model features influence predictions. These tools therefore address complementary aspects of trustworthiness and predictive generalization, but none by itself establishes whether an estimator of prediction reliability remains accurate after transfer to a different biological context. As a result, we still lack principled approaches for measuring reliability transfer, identifying suitable reliability-transfer sources, or understanding why some source-target pairs transfer reliability successfully whereas others do not. Addressing these questions requires a framework capable of characterizing and comparing the biological and contextual signals that models use when determining whether predictions can be trusted across diverse contexts.

Here, we introduce scPerturbTrust, a framework for understanding and improving reliability transfer across single-cell foundation models. We hypothesized that reliability transfers more successfully when models rely on similar biological and contextual signals to determine whether their predictions can be trusted. For example, reliability may depend on different combinations of network connectivity, cellular localization, target exposure, and contextual information across biological settings. As a result, two contexts can achieve similar predictive performance while differing substantially in how prediction reliability is assessed. To test this hypothesis, we introduce reliability-attribution fingerprints, compact signatures designed to characterize how reliability is determined within a biological context. Unlike conventional attribution analyses ^10, 11^, which focus on explaining individual predictions, these fingerprints are specifically designed to compare reliability behavior across contexts. Using strictly cross-fitted SHAP attributions, we construct fingerprints that can be compared across datasets without using target outcomes for explanation. Across ten model–dataset systems, we show that reliability transfer is a distinct form of generalization that cannot be inferred from predictive performance alone. We further demonstrate that contexts with similar reliability-attribution fingerprints tend to transfer reliability more successfully, revealing structured relationships between reliability behavior and transferability. Together, these findings establish reliability transfer as a distinct dimension of foundation-model generalization and introduce reliability-attribution fingerprints as a practical framework for characterizing transferable reliability structure. More broadly, our results suggest that the utility of foundation models depends not only on transferring predictive knowledge across biological contexts, but also on transferring a reliable understanding of when that knowledge can be trusted.

### Reliability varies across biological contexts

To establish whether prediction reliability is uniform within a model–data system or varies across perturbation queries, we first analyzed the Norman Perturb-seq benchmark using the GEARS processing framework and scGPT perturbation prediction pipeline (Fig. 1A; Methods). The benchmark comprised 91,205 cells and 284 perturbation conditions, including 116 held-out test perturbations spanning a range of target-gene exposure histories. Prediction reliability varied substantially across perturbations evaluated using the same prediction workflow. Representative held-out perturbations illustrated that scGPT closely reproduced observed transcriptional responses for some queries but deviated considerably for others (Fig. 1B). Thus, average performance at the model–dataset level obscured substantial differences in reliability among individual perturbations. ^12^ ^1^ ^2^

**Fig. 1.**
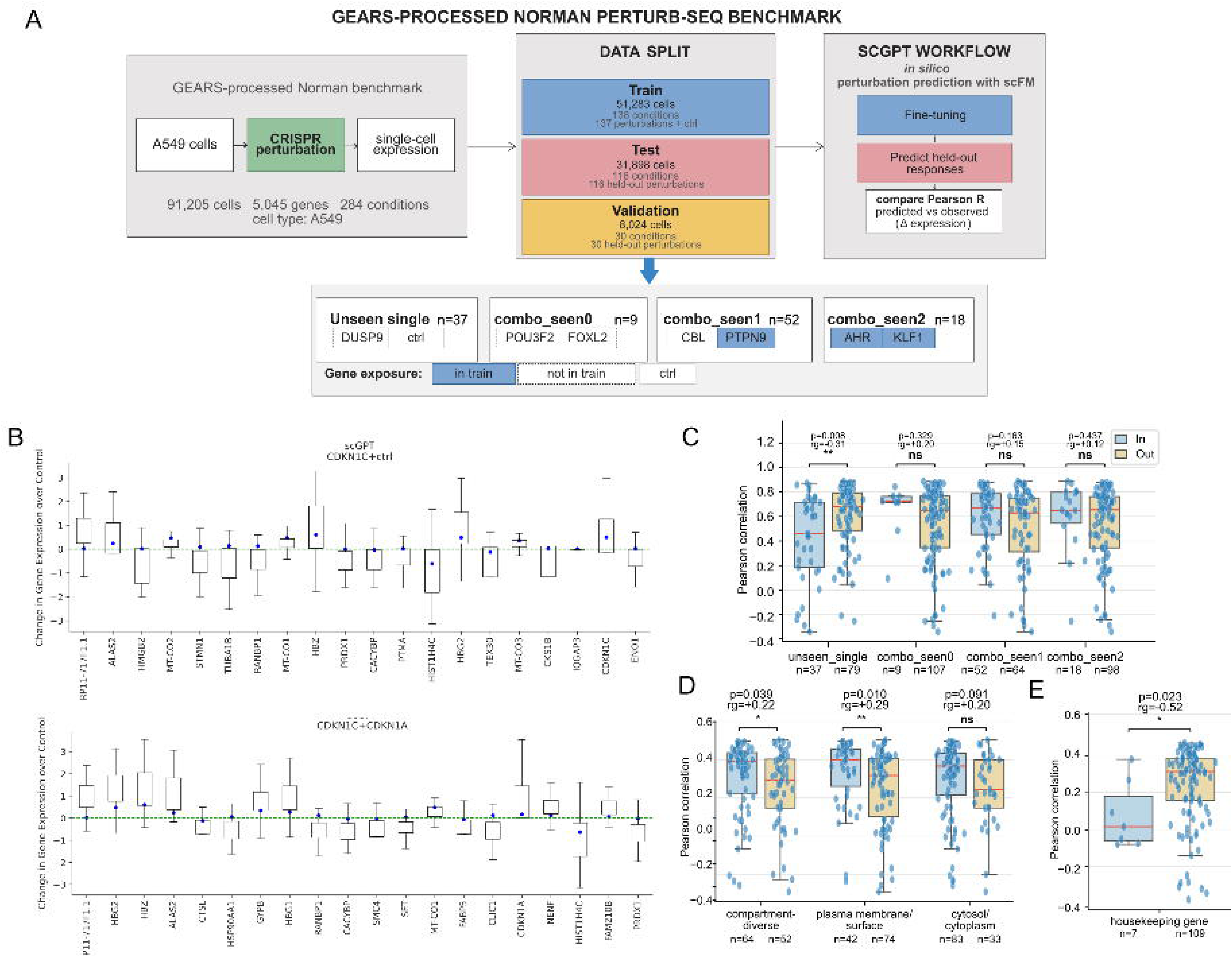
**(A)** Overview of the GEARS-processed Norman Perturb-seq benchmark, predefined data partition, scGPT prediction workflow and held-out perturbation exposure categories. **(B)** Representative scGPT predictions for held-out Norman perturbations. Boxplots show observed single-cell expression changes relative to the control mean; blue points show predicted condition-level changes, and the green dashed line denotes no change from control. **(C-E)** Query-level Pearson correlations between predicted and observed delta-expression profiles, stratified by **(C)** GEARS exposure category, **(D)** selected cellular-component features and **(E)** housekeeping-gene membership. “In” and “Out” denote feature-positive and feature-negative queries, respectively. Points represent perturbation queries; boxes show medians and interquartile ranges. P values are from two-sided Wilcoxon rank-sum tests, and *r_g_* denotes the rank-biserial effect size \**p*<0.05,\*\**p*<0.01 ns, not significant.

To investigate potential sources of this variation, we stratified perturbations according to prior target exposure and selected biological annotations. Unseen single-gene perturbations exhibited lower performance than other held-out perturbations (two-sided Wilcoxon P = 0.008, rank-biserial effect size *r_g_*= 0.31; Fig. 1C). In contrast, double perturbations showed no consistent performance differences based on target-gene exposure history (all P ≥ 0.163). Reliability also varied across selected biological categories (Fig. 1D-E). Perturbations involving compartment-diverse genes or plasma-membrane and cell-surface genes showed higher performance (P = 0.039, *r_g_*=0.22; P = 0.010, *r_g_*=0.29), whereas perturbations involving housekeeping genes tended to show lower performance; P show lower performance (*r_g_*=0.52, nominal P < 0.05), although the latter comparison involved only seven held-out queries and should therefore be interpreted cautiously. Together, these results demonstrate that prediction reliability is not a uniform property of a perturbation-prediction system. Instead, substantial reliability differences emerge across perturbations even within a single biological context, suggesting that reliability depends on characteristics of both the query and its underlying biological setting. This heterogeneity motivates the central question of this study: whether such reliability patterns can be transferred across biological contexts and, if so, what determines successful reliability transfer.

### Prediction transfer does not imply reliability transfer

Having established that reliability varies substantially within individual model–dataset systems, we next asked whether reliability learned in one biological context could be transferred to another. Pairwise source-to-target reliability classification ^8^ revealed substantial heterogeneity in transfer performance, spanning a wide range of ROC AUCs across classifier families (Fig. 2A-B; Supplementary Fig. 1A-E). For the same target system, transfer performance often differed markedly depending on the source system used to train the reliability estimator. Despite differences in classifier architecture, similar transfer patterns were observed across methods, with source-target pairs that transferred well in one classifier generally showing strong transfer in others, whereas poorly transferring pairs remained consistently weak. To determine whether transferability could be explained by biological information associated with the query itself, we compared two reliability estimators: an exposure/context model (M2) and a safeguarded variant (M3) that incorporated biological annotation features while reducing opportunities for information leakage (Methods). Both estimators were evaluated under increasingly stringent leave-group-out settings, including gene-purged analyses that excluded both held-out contexts and their constituent query genes from training (Fig. 2C). Performance differences between M2 and M3 varied across datasets, model families, and model–dataset systems.

**Fig. 2:**
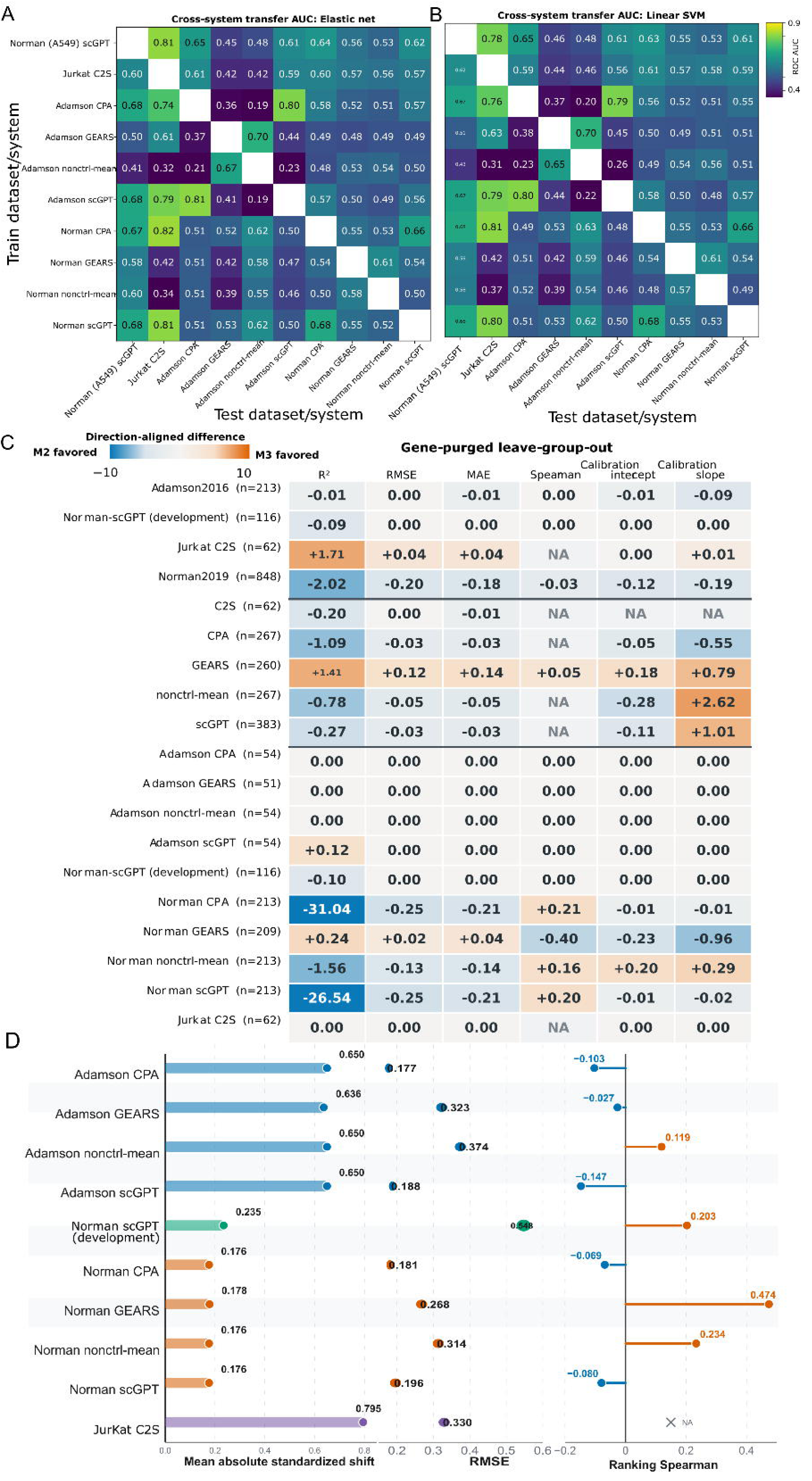
Reliability transfer defines a distinct form of generalization. Pairwise source-to-target ROC AUC for binary reliability classification using (A) elastic-net logistic regression, (B) linear support-vector machine. (C) Direction-aligned performance difference between safeguarded M3 and M2 after withholding a source dataset, model family or model–dataset system. Evaluation excludes the held-out group and training queries containing genes represented in that group. Positive values (red) favor M3, negative values (blue) favor M2, and hatched cells were not evaluable. (D) Decomposition of standard leave-system-out transfer into covariate shift, prediction error, query ranking and calibration error. Colors denote source datasets.

We next decomposed leave-system-out transfer behavior into contributions from covariate shift, prediction error, query ranking, and calibration effects ^6, 13^ (Fig. 2D; Methods). The magnitude of these components differed across held-out target systems. Covariate shift ranged from 0.176–0.235 among the Norman targets to 0.795 for Jurkat C2S, while query-ranking correlations ranged from −0.147 for Adamson nonctrl-mean to 0.474 for Norman GEARS. Prediction error (RMSE) and calibration metrics also varied among target systems. Thus, different target systems exhibited distinct transfer profiles across the four decomposition components. Together, these results show that reliability transfer varies markedly across source-target pairs and is not determined solely by dataset identity, model family, or individual reliability estimators. This variability motivated our subsequent search for transferable reliability structure that could explain and predict successful transfer across biological contexts.

### Shared and context-specific biological signals influence prediction reliability

To investigate whether prediction reliability is associated with common biological features across systems, we quantified adjusted associations between query-level performance and biological annotations ^14, 15^ while controlling for exposure and contextual effects (Fig. 3A; Methods). Several biological features exhibited recurring patterns across subsets of systems. For example, plasma-membrane and cell-surface annotations were frequently associated with higher reliability, whereas mitochondrial features were often associated with lower reliability. However, the magnitude and significance of these effects varied substantially among model–dataset systems. Other features, including transcription-factor targets, housekeeping genes, and regulatory categories, exhibited stronger context dependence, with effect sizes differing markedly across systems. These results suggest that prediction reliability reflects a combination of shared and system-specific biological signals rather than a universal set of determinants.

**Figure 3.**
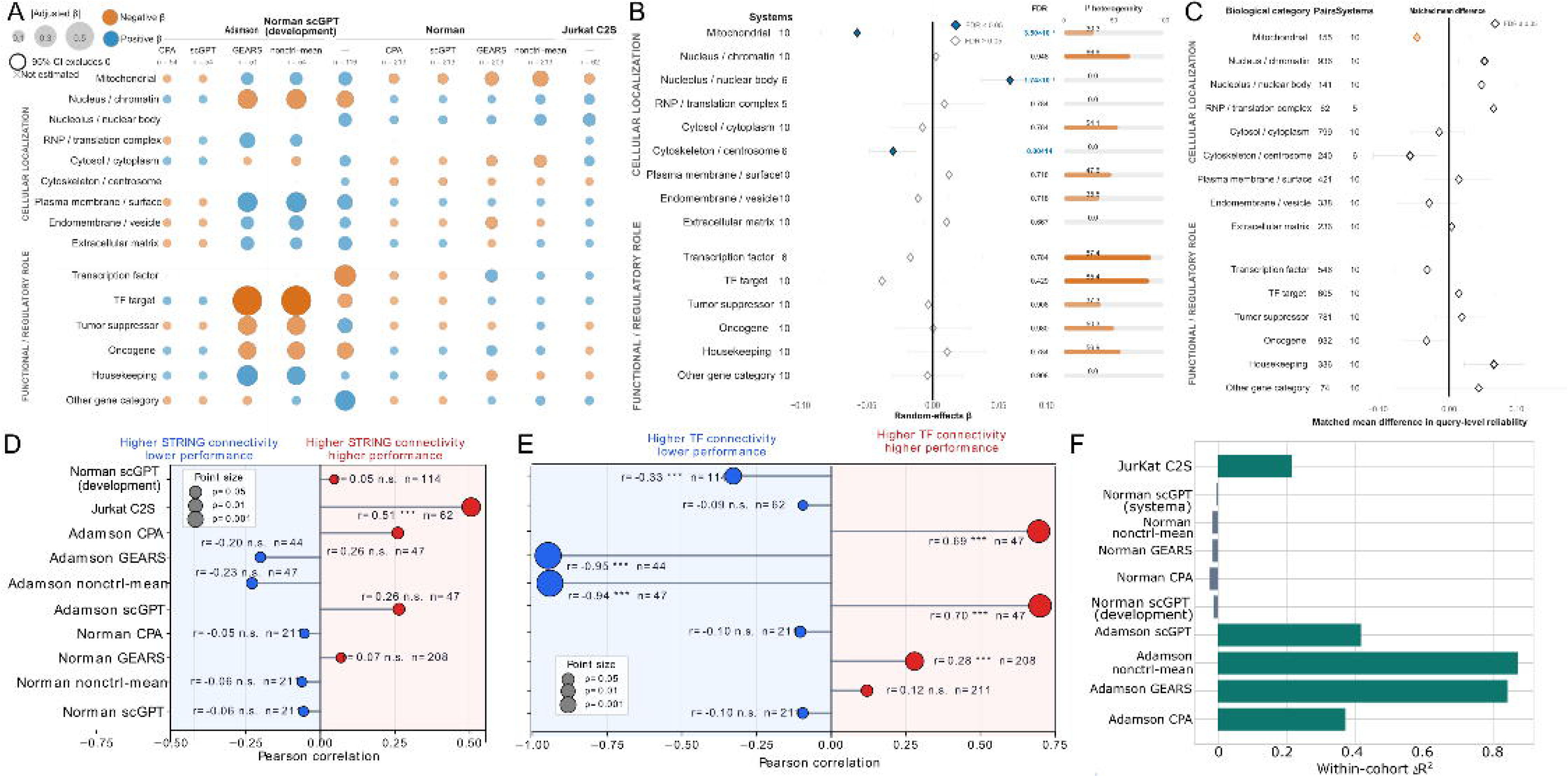
Reliability is governed by context-specific signals. (A) System-specific adjusted associations between biological features and query-level prediction performance. Circle area represents the absolute adjusted coefficient, color denotes its direction, and outlined circles indicate estimates whose 95% confidence intervals exclude zero. Crosses indicate features that could not be estimated in a system. (B) Random-effects meta-analysis of adjusted biological-feature associations. Diamonds and horizontal lines show pooled coefficients and 95% confidence intervals. Filled diamonds denote FDR. Adjacent columns report the number of evaluable systems, FDR-adjusted significance and heterogeneity. (C) Matched differences in query-level performance between feature-positive and feature-negative perturbations. Diamonds and horizontal lines show matched mean differences and bootstrap 95% confidence intervals. Filled diamonds denote FDR. The numbers of matched pairs and contributing systems are shown. (D, E) Within-system Pearson correlations between prediction performance and (D) high-confidence STRING connectivity or (E) CollecTRI TF-target connectivity. Red indicates positive and blue indicates negative correlations; point size reflects nominal statistical significance. (F) System-specific difference in out-of-fold between safeguarded M3 and M2. Positive values favor M3.

To identify signals that generalized across systems, we performed a random-effects meta-analysis of the adjusted feature associations (Fig. 3B; Methods). Nucleolus/nuclear-body localization exhibited a significant positive association with reliability (β ≈ 0.06, FDR = 1.74 × 10⁻□), whereas mitochondrial localization (β ≈ −0.06, FDR = 3.50 × 10⁻□) and cytoskeleton/centrosome localization (β ≈ −0.04, FDR = 0.004) showed significant negative associations. Most remaining features were not significant after multiple-testing correction and several exhibited substantial between-system heterogeneity, indicating that shared biological effects coexist with substantial system-specific variation.

Confounder-matched analyses yielded broadly similar trends (Fig. 3C; Methods). Associations detected in the meta-analysis were generally preserved after matching feature-positive and feature-negative perturbations with similar characteristics, although effect sizes were typically smaller. These results support the robustness of several shared biological associations while highlighting their modest magnitude at the perturbation level.

We next examined whether network structure was associated with reliability. Correlations between reliability and STRING connectivity ^16^ differed substantially across systems (Fig. 3D). For example, reliability was positively associated with connectivity in Jurkat C2S (r = 0.51, P < 0.001) but weakly negative in Adamson GEARS (r = −0.23) and Adamson nonctrl-mean (r = −0.20). Similar context dependence was observed for CollecTRI transcription factor–target connectivity ^17^ (Fig. 3E), ranging from strongly negative correlations in Adamson GEARS (r = −0.95, P < 0.001) and Adamson nonctrl-mean (r = −0.94, P < 0.001) to strongly positive correlations in Adamson scGPT (r = 0.70, P < 0.001) and Jurkat C2S (r = 0.69, P < 0.001).

To assess whether these biological signals improve reliability estimation, we compared an exposure/context model (M2) with a safeguarded variant (M3) that incorporated biological annotation features while reducing opportunities for information leakage (Fig. 3F; Methods). The contribution of biological annotations varied across systems. M3 substantially increased within-cohort explanatory power for Adamson nonctrl-mean (ΔR² ≈ 0.87), Adamson GEARS (ΔR² ≈ 0.84), Adamson scGPT (ΔR² ≈ 0.41), and Adamson CPA (ΔR² ≈ 0.37), while providing comparatively little improvement for several Norman systems.

Together, these results indicate that prediction reliability is associated with both shared and context-specific biological signals. A limited set of biological features shows reproducible associations across systems, but the strength, direction, and predictive utility of those signals vary substantially among model–dataset systems. This structured variation motivated our subsequent search for a representation capable of comparing reliability behavior across biological contexts.

### Reliability-attribution fingerprints capture how reliability is determined across contexts

We now sought a representation that could characterize reliability behavior within a context and enable comparisons across contexts. To this end, we computed strictly out-of-fold SHAP values ^10, 11^ for fitted reliability estimators and aggregated feature contributions into eight predefined domains: system context, regulatory connectivity, network context, cellular localization, perturbation complexity, expression support, gene category, and model exposure (Fig. 4A; Methods). The resulting domain-level attribution profiles constituted reliability-attribution fingerprints. Signed fingerprints summarized directional effects, whereas magnitude fingerprints summarized domain importance irrespective of direction.

**Fig. 4:**
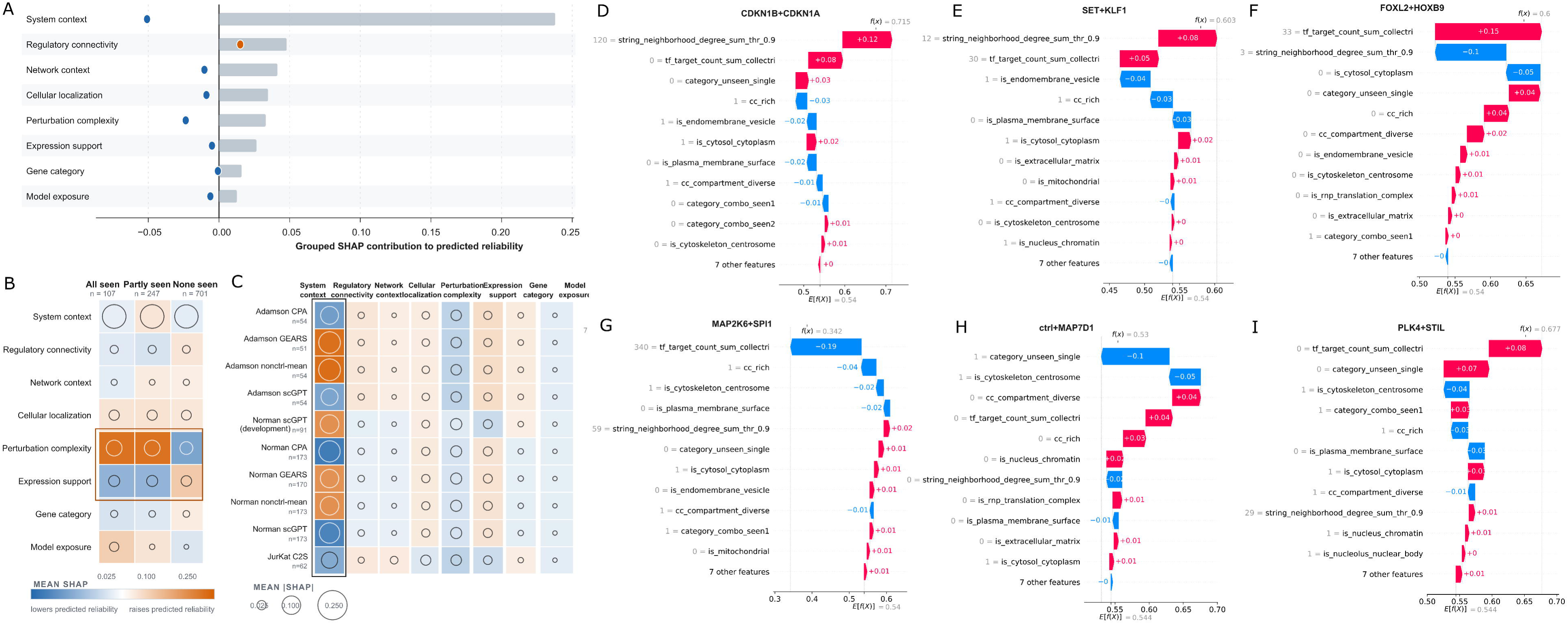
Reliability-attribution fingerprints capture how reliability is determined in each context. **(A)** Construction of reliability-attribution fingerprints using cross-fitted SHAP explanations of reliability estimators. Individual attribution values are aggregated to generate context-level signatures of reliability behavior. **(B)** Attribution fingerprints stratified by perturbation exposure category, illustrating differences in reliability structure within a system. **(C)** Reliability-attribution fingerprints across model–dataset systems. Distinct attribution patterns indicate that systems rely on different combinations of exposure, contextual, and biological signals when assessing prediction reliability. **(D–I)** Representative SHAP waterfall plots illustrating perturbation-specific reliability explanations. These examples demonstrate how individual biological and contextual features contribute to reliability estimation and collectively motivate the use of attribution fingerprints as context-level representations of reliability behavior.

Across 3,920 out-of-fold predictions, system context showed the largest average attribution magnitude (mean absolute grouped SHAP = 0.238), with regulatory connectivity, network context, cellular localization, perturbation complexity, and exposure-related domains contributing smaller but distinguishable effects (Fig. 4A). Reliability estimates therefore reflected combinations of contextual, exposure-related, and biological information rather than any single dominant feature class.

Reliability-attribution fingerprints varied both by target exposure and by model–dataset system (Fig. 4B,C). Perturbation complexity contributed positively when all or some target genes had previously been observed during perturbation-model training (signed mean SHAP = 0.096 and 0.095, respectively), but negatively when no targets had been observed (−0.060). Conversely, expression support contributed negatively in the all-seen and partly-seen groups (−0.048 in both) and positively in the none-seen group (0.028). Distinct system-level fingerprints were similarly evident across Adamson, Norman, and Jurkat analyses, indicating that different systems relied on different combinations of biological and contextual signals when assessing prediction reliability.

Representative SHAP explanations ^10, 11^ illustrated how these fingerprints operate at the level of individual predictions (Fig. 4D–I). Perturbations with similar baseline reliability estimates nonetheless received substantially different reliability predictions because regulatory connectivity, target exposure, cellular localization, and other contextual features contributed differently to the final estimate. In the regulatory-connectivity examples (Fig. 4D–G), increasing connectivity was generally associated with lower estimated reliability, whereas the exposure-matched examples (Fig. 4H–I) highlighted the contribution of target novelty. These examples demonstrate how reliability-attribution fingerprints decompose reliability estimates into interpretable biological and contextual contributions. Several attribution effects were also conditional on interactions among biological and exposure-related features (Supplementary Fig. 3A,B; Methods), indicating that reliability estimation was context dependent rather than purely additive.

Together, these analyses establish reliability-attribution fingerprints as context-level representations of reliability behavior. Rather than identifying causal determinants of predictability, these fingerprints characterize how fitted reliability estimators combine biological and contextual information when determining whether predictions should be trusted. We next tested whether similarity in these fingerprints predicts successful reliability transfer across biological contexts.

### Shared reliability structure predicts successful transfer

To test whether successful reliability transfer depends on shared reliability structure, we compared transfer performance with similarity in reliability-attribution fingerprints derived from the source and target systems (Fig. 5A-C). Across targets, source systems with more similar attribution fingerprints generally transferred reliability more successfully than dissimilar sources. This relationship was most evident for fingerprint magnitudes, where the within-target correlation between fingerprint distance and transfer performance was consistently negative (mean Spearman ρ ≈ −0.43), indicating that sources with more similar reliability-attribution structure tended to achieve higher transfer performance (Fig. 5A). By contrast, signed-cosine and signed-correlation representations produced positive correlations, suggesting substantially weaker alignment with transferability.

**Figure 5:**
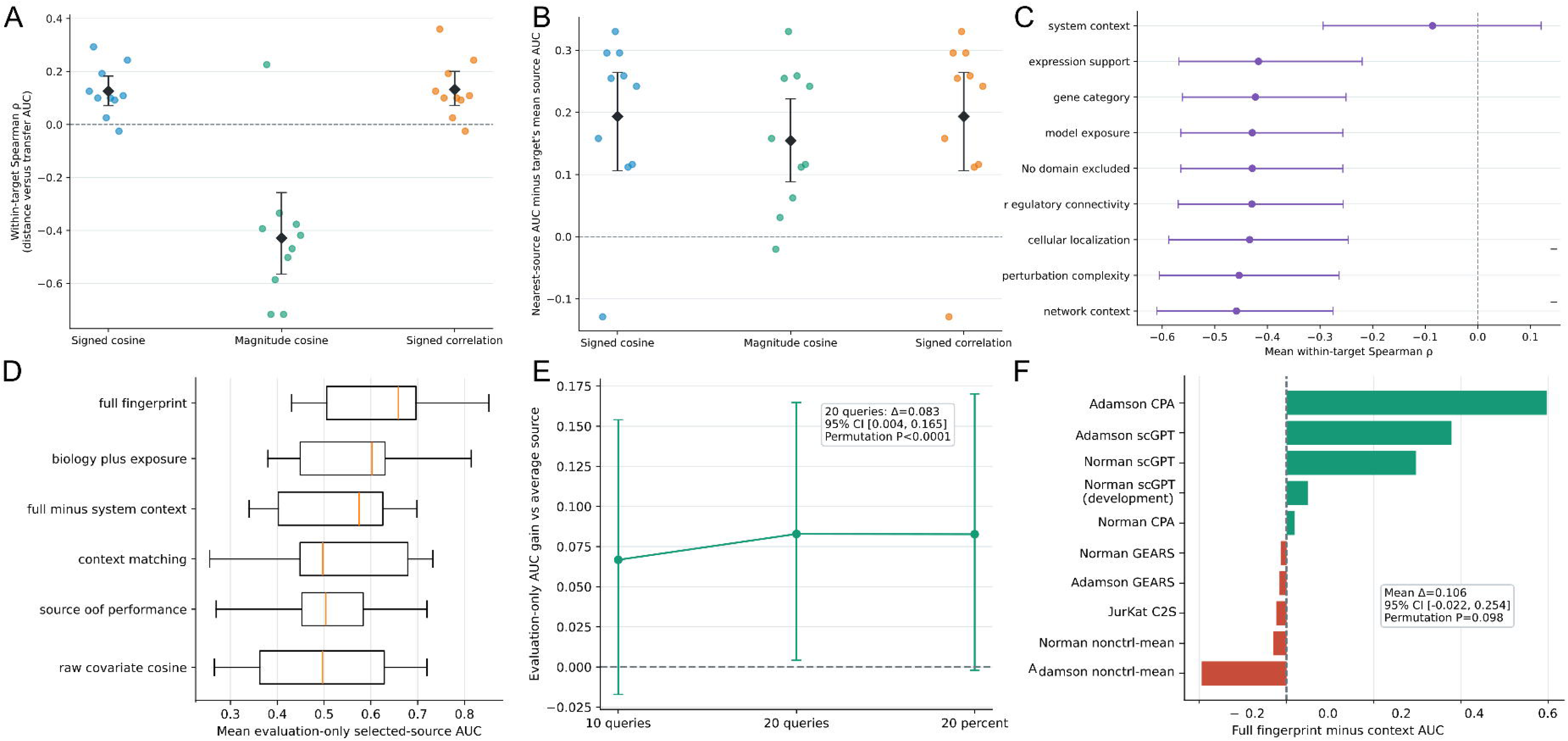
Similarity in reliability-attribution fingerprints predicts transfer success. **(A)** Within-target Spearman correlations between fingerprint similarity and source-to-target reliability-transfer performance using signed cosine similarity, magnitude cosine similarity, and signed correlation. Each point corresponds to a target system; diamonds indicate means and error bars show 95% confidence intervals. **(B)** Improvement in transfer performance obtained by selecting the fingerprint-nearest source relative to the target-average source. Positive values indicate improved reliability transfer. **(c)** Within-target fingerprint-transfer correlation after excluding individual attribution domains. More negative values indicate stronger agreement between fingerprint similarity and transfer performance. The dashed vertical line indicates no association. **(D)** Mean transfer performance obtained using different source-selection strategies, including full fingerprints, biology-plus-exposure fingerprints, fingerprints excluding system context, context matching, source-performance heuristics, and raw covariate similarity. **(E)** Evaluation-only source-selection performance under practical deployment settings. Error bars represent 95% confidence intervals. Annotation indicates mean AUC improvement relative to the average source and the corresponding permutation-test significance. **(F)** Change in source-selection performance after removing system-context information from the fingerprint representation. Positive values indicate that system context improved source identification. Error bars and annotation summarize the average effect across systems.

This relationship also translated into source-selection performance. Selecting the most similar source according to fingerprint similarity yielded mean transfer gains of approximately 0.15 AUC over the target-average source across multiple targets (Fig. 5B). Similar improvements were observed for both signed-correlation and signed-cosine measures, whereas magnitude-based fingerprints consistently identified compatible transfer sources. These results indicate that reliability-attribution fingerprints contain information relevant to transferability that is not captured by source identity alone.

To determine which fingerprint components contributed most strongly to this relationship, we systematically excluded individual attribution domains and recalculated within-target fingerprint-transfer correlations (Fig. 5C). Removing system-context information produced the largest reduction in performance, shifting the mean within-target correlation from approximately −0.43 toward zero. Excluding expression support, gene category, model exposure, regulatory connectivity, cellular localization, perturbation complexity, or network context also weakened the relationship, although to a lesser extent. Thus, transferability was associated with multiple fingerprint domains, with system context contributing the largest single component.

We next evaluated whether fingerprint similarity could be used directly to identify transferable reliability sources. Across all systems, the full fingerprint outperformed context matching, source-performance heuristics, and raw covariate similarity as a source-selection strategy (Fig. 5D). Evaluation-only source selection produced mean selected-source AUCs near 0.64, whereas raw covariate similarity and source-performance heuristics achieved lower values near 0.50–0.55. Fingerprint-guided selection remained effective across practical deployment settings, improving transfer AUC by approximately 0.08 over the average source when evaluation was limited to 20 held-out queries and producing similar gains when selecting 20% of available queries (Fig. 5E).

Finally, the contribution of system-context information varied across model–dataset systems (Fig. 5F). Removing this domain reduced source-selection performance most strongly for Adamson CPA, Adamson scGPT, and Norman scGPT, whereas several Norman and Jurkat systems showed little change. These results suggest that although multiple biological and contextual domains contribute to transferability, their importance differs among systems.

Together, these analyses support the central hypothesis of this study: contexts exhibiting similar reliability-attribution fingerprints transfer reliability more successfully. Reliability-attribution fingerprints therefore provide a quantitative representation of transfer compatibility and reveal previously unrecognized structure underlying reliability transfer across biological contexts.

### Reliability transfer can alter perturbation rankings

Having shown that reliability-attribution fingerprints can identify compatible transfer sources, we next examined whether the choice of reliability source could influence perturbation prioritization ^18^. Using Norman scGPT as a representative target system, we compared perturbation rankings generated by reliability estimators transferred from the fingerprint-nearest source (Norman CPA; transfer AUC = 0.803) and the fingerprint-most-distant source (Norman scGPT; transfer AUC = 0.601) (Fig. 6A; Methods). Although both estimators were applied to the same set of 213 perturbation predictions, several perturbations changed substantially in rank between the two transfers. Consistent with these ranking differences, the fingerprint-nearest source produced higher observed performance among prioritized perturbations. For the top 10 ranked perturbations, the mean observed Pearson delta correlation was 0.189 for the nearest-source transfer compared with 0.168 for the distant-source transfer, with 10 of 10 versus 8 of 10 perturbations exceeding the target-wide median performance. At corresponding mean observed performances were 0.184 and 0.152, with 20 of 20 versus 15 of 20 perturbations exceeding the median. Because this analysis uses the full target dataset for fingerprint calculation and summarization, it should be interpreted as an illustrative case study rather than independent held-out validation.

**Figure 6:**
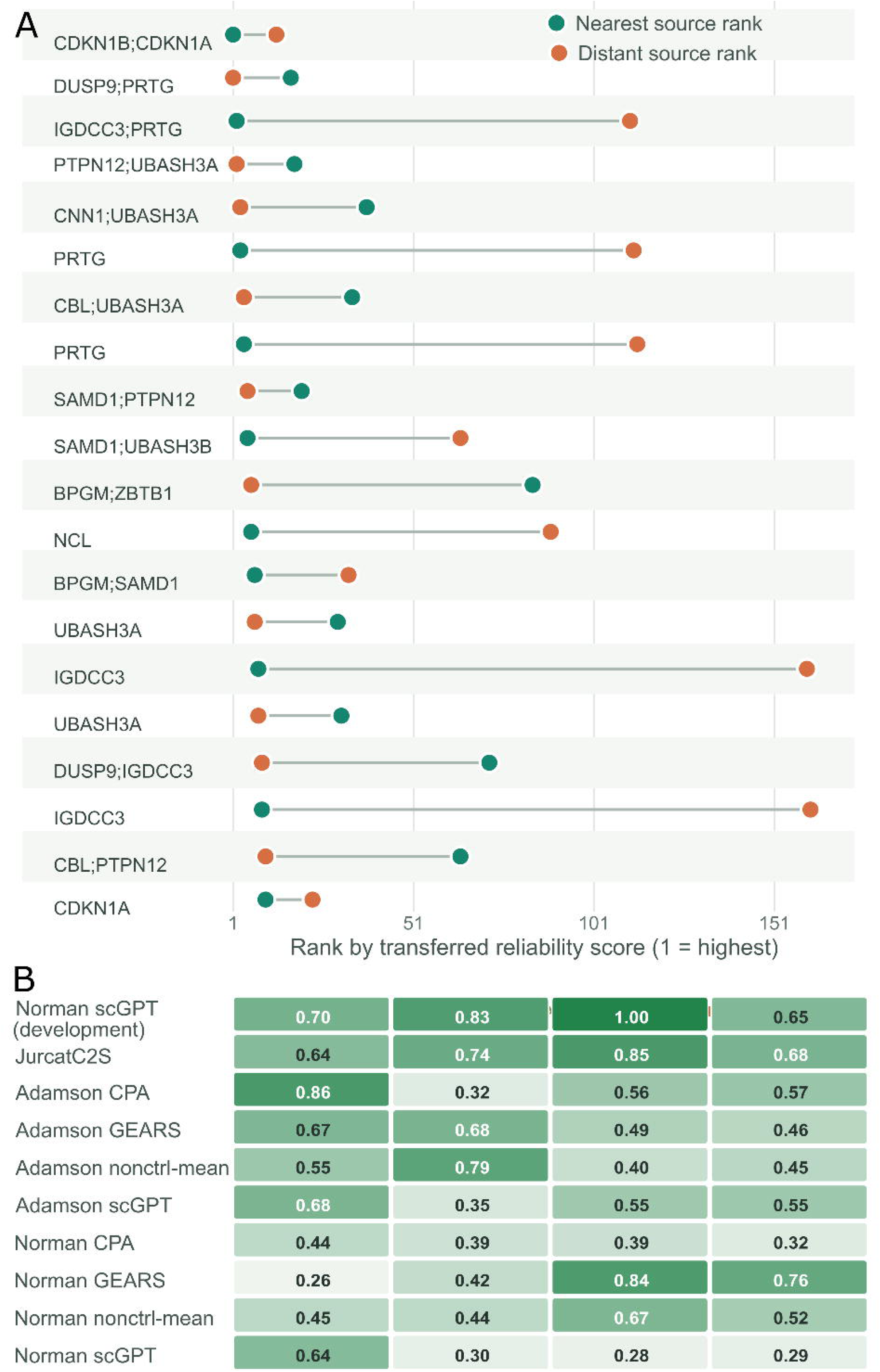
Source-specific reliability estimates alter perturbation rankings. **A** For Norman scGPT, each row is a perturbation appearing in at least one source’s top-10 list. Points show its rank among all 213 target perturbations based on the transferred reliability score: green, fingerprint-nearest source; orange, fingerprint-distant source. Rank 1 is the highest priority; connecting lines pair the same perturbation across sources. **B,** Heat map of results across target systems and four source-selection strategies: fingerprint-nearest, context-matched, average source, and random source. Darker green indicates larger values.

We next assessed whether such prioritization differences generalized across systems using constituent-gene-disjoint calibration and evaluation splits spanning ten target systems (Fig. 6B; Supplementary Fig. 3; Methods). At a calibration budget of 20 queries, fingerprint-nearest source selection achieved a mean precision@10 of 0.587 and mean observed performance@10 of 0.352. The corresponding values were 0.526 and 0.367 for context matching, 0.603 and 0.395 for average-source scoring, and 0.526 and 0.360 for random-source selection. Similar patterns were observed at *K=20*, where fingerprint-nearest selection achieved a precision@20 of 0.513 and a mean observed performance@20 of 0.344, compared with 0.430 and 0.341 for context matching, 0.573 and 0.397 for average-source scoring, and 0.479 and 0.351 for random-source selection. However, fingerprint-nearest source selection did not significantly outperform any comparator for either precision or mean observed performance at either ranking threshold (all paired exact-test *p* ≥ 0.232.

Together, these analyses indicate that transferred reliability estimates can influence perturbation prioritization and, in selected targets, may enrich highly performing perturbations among the top-ranked candidates. However, the population-level analyses did not demonstrate a consistent prioritization advantage for fingerprint-nearest source selection over context matching, source averaging, or random-source baselines. These findings therefore support a practical consequence of reliability transfer while suggesting that the downstream benefits of attribution-guided source selection remain context dependent.

## Discussions

Single-cell foundation models are increasingly evaluated by their ability to transfer predictive performance across biological contexts ^1, 2, 4^. Our results suggest that a second form of generalization should also be considered: reliability transfer, the ability to transfer an accurate assessment of when predictions can be trusted. Across ten model–dataset systems, reliability transfer varied substantially among source-target pairs and could not be inferred from predictive performance alone. Thus, successful prediction transfer does not necessarily imply successful reliability transfer.

To investigate this variability, we examined the biological and contextual signals associated with prediction reliability. Some signals, including selected cellular-localization features, showed reproducible associations across systems, whereas others varied substantially in direction and magnitude. Network connectivity and biological annotations likewise contributed differently across contexts. These findings suggest that prediction reliability is shaped by a combination of shared and context-dependent signals rather than a single universal determinant.

Motivated by this observation, we introduced reliability-attribution fingerprints, context-level summaries of how reliability estimators combine biological and contextual information when assigning confidence ^10, 19^. Attribution-magnitude fingerprints were associated with source-ranking performance within the observed candidate set and identified transfer sources that achieved higher transfer AUC than the average available source. However, these relationships weakened when candidate sources from the same dataset family were excluded and depended substantially on system-context information. Accordingly, our results support attribution-guided source matching as a proof of concept rather than a general solution for reliability transfer across unrelated biological settings.

The perturbation-prioritization analyses illustrate the practical consequences of this problem. Different transferred reliability estimators produced different perturbation rankings despite identical perturbation predictions, showing that reliability transfer can influence downstream evaluation decisions. However, population-level analyses did not establish a consistent prioritization advantage for fingerprint-guided source selection over context matching, source averaging, or random selection. Thus, although transferred reliability estimates can affect prioritization, the practical benefit of attribution-guided source selection remains context dependent.

Several limitations should be noted. First, the analyses were restricted to perturbation-prediction systems and may not generalize to all biological foundation models. Second, reliability-attribution fingerprints describe how fitted reliability estimators generate confidence estimates and should not be interpreted as causal determinants of predictability ^19^. Third, the transfer relationships identified here were evaluated within a finite collection of datasets and models. Finally, prioritization outcomes were assessed using retrospective prediction performance rather than prospective experimental outcomes.

Despite these limitations, our findings identify reliability transfer as a distinct aspect of foundation-model generalization. Prediction transfer addresses whether knowledge learned in one biological context remains useful in another; reliability transfer addresses whether the model can accurately assess the limits of that knowledge after transfer. As foundation models become increasingly integrated into biological discovery pipelines, understanding and evaluating reliability transfer may become as important as evaluating predictive performance itself. Reliability-attribution fingerprints provide a first framework for studying this problem and for characterizing transferable reliability structure across biological contexts.

## Methods

### Study systems and definition of reliability

We analyzed ten perturbation-prediction systems comprising four prediction methods evaluated on the Adamson Perturb-seq dataset ^20^, four methods evaluated on the Norman Perturb-seq dataset ^12^, the original Norman scGPT development system, and the Jurkat C2S benchmark ^21^. For each perturbation query *q*, prediction reliability was defined as the Pearson correlation between predicted and observed perturbation-induced expression changes,

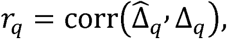

Where Δ_q_and Δ_q_denote the predicted and observed perturbation-induced expression vectors, respectively. When multiple Systema seeds were available, reliability was summarized using the median correlation across seeds ^22^.

### Query-level biological and contextual features

Perturbed genes were annotated using nine cellular-component categories derived from the precomputed binary feature matrix and six regulatory or functional categories: transcription factor, TF target, tumor suppressor, oncogene, housekeeping gene, and other. These annotations were compiled from curated transcription-factor catalogs ^15^, TRRUST ^23^, CancerMine ^24^, and housekeeping-gene reference sets ^25, 26^. For multi-gene perturbations, a feature was considered present if any constituent target gene carried the annotation.

Additional query-level variables included perturbation size, prior target exposure, expression support, STRING protein–protein interaction connectivity ^16^, CollecTRI transcription factor–target connectivity ^17^, cellular-component richness, and system-context indicators.

Prior target exposure was defined as the fraction of constituent perturbation genes previously observed during training of the underlying perturbation-prediction model,

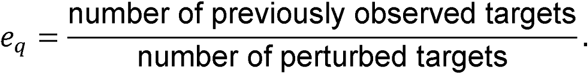

### Reliability models

Three classes of reliability estimators were considered.

The baseline model, *M*_0_, assigned every query the mean reliability observed within the corresponding training partition.

The exposure/context model, *M*_2_, incorporated only pre-outcome variables available without biological annotations, including perturbation size, target exposure, expression-support measures, and contextual features.

The biology-augmented model, *M*_3_, extended *M*_2_by incorporating biological annotation features, including cellular localization, network connectivity, regulatory annotations, and curated gene-category labels.

To reduce opportunities for overfitting and information leakage, *M*_3_was retained only when biological annotation features improved training-only validation performance. Otherwise, the fitted *M*_2_model was used. This safeguarded procedure ensured that biological features were incorporated only when they provided demonstrable predictive value.

### Constituent-gene-disjoint validation

Primary within-system evaluation used five outer folds repeated five times. Folds were constructed at the constituent-gene level ^27^. A perturbation query was eligible for an outer test fold only when all constituent target genes were absent from the corresponding training partition.

Hyperparameter tuning, feature selection, preprocessing, standardization, and imputation were performed exclusively within the training data of each fold. Inner cross-validation used the same constituent-gene-disjoint constraint ^27, 28^.

All reported reliability predictions were generated from out-of-fold evaluations.

### Biological determinants of reliability

To quantify associations between biological features and prediction reliability, we fitted a separate ordinary least-squares model for each binary feature within each prediction system:

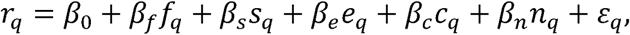

where:

- *r_q_* is query reliability,
- *f_q_* denotes features membership,
- *s_q_* is perturbation size,
- *e_q_* is prior target exposure,
- *c_q_* is cellular-component richness,
- *n_q_* is STRING connectivity.

Continuous covariates were standardized within system. Missing values were median-imputed where applicable.

Models were fitted only when:

1. at least 25 queries were available,
2. both feature states were represented,
3. the smaller feature group contained at least four queries.

The adjusted coefficient *β_f_* was interpreted as the reliability difference between feature-positive and feature-negative perturbations after adjustment for covariates.

### Meta-analysis and confounder-matched analyses

System-specific coefficients were combined using random-effects meta-analysis.

To assess robustness, feature-positive perturbations were matched to feature-negative perturbations with similar values of exposure, perturbation size, and annotation richness. Reliability differences were calculated within matched pairs and summarized using bootstrap confidence intervals.

### Connectivity analyses

Within each prediction system, Pearson correlations were calculated between query-level reliability and:

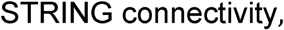

And

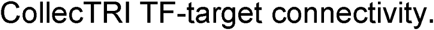

These analyses assessed whether network structure was associated with prediction reliability.

### Cross-fitted SHAP attribution

SHAP values were computed exclusively for queries withheld from reliability-model fitting.

For each outer-validation partition:

1. a reliability estimator was fitted using training queries,
2. SHAP values were computed for outer-test queries,
3. the test-query outcome was never used to fit the model generating its explanation.

TreeSHAP was used for nonlinear estimators ^11, 29^. Exact SHAP values were computed for linear estimators ^10^. SHAP additivity was verified by confirming

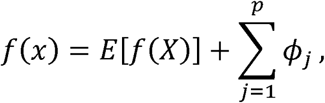

where:

- *f(x)* is the reliability prediction,
- *E[f(X)]* is the fold-specific expected prediction,
- *Фj* is the SHAP contribution of feature *j*,

### Reliability-attribution fingerprints

Individual features were grouped into eight predefined attribution domains:

1. system context,
2. regulatory connectivity,
3. network context,
4. cellular localization,
5. perturbation complexity,
6. expression support,
7. gene category,
8. model exposure.

For a prediction *i*, grouped SHAP values were computed as

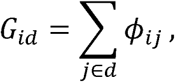

Where *d* denotes an attribution domain.

For each context, two fingerprint representations were defined.

The signed fingerprint:

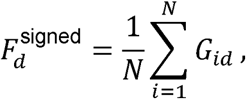

summarized domain importance independent of direction. These vectors constituted the reliability

The magnitude fingerprint:

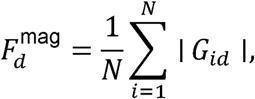

summarized domain importance independent of direction.

These vectors constituted the reliability-attribution fingerprints.

### Exposure- and system-specific fingerprints

Fingerprints were summarized separately according to target exposure category:

- all targets observed,
- partially observed targets,
- no observed targets.

System-level fingerprints were calculated independently for each of the ten prediction systems.

Within each stratum, signed mean SHAP quantified directionality and mean absolute SHAP quantified attribution magnitude.

### Pairwise reliability transfer

For each ordered source-target pair, a binary reliability classifier was trained in the source system and evaluated in the target system.

Elastic-net logistic regression and linear support-vector machines ^30, 31^ served as primary classifiers. Additional classifier families were evaluated as sensitivity analyses.

Transfer performance was summarized using target-system ROC AUC. Source-to-self comparisons were excluded.

### Leave-group-out reliability transfer

Transfer analyses were performed under three withholding schemes:

1. source dataset,
2. prediction-model family,
3. individual model–dataset system.

Under standard leave-group-out evaluation, all observations from the held-out group were excluded from training.

Under gene-purged evaluation, all training queries containing any constituent gene represented in the held-out group were additionally removed.

Both *M*_2_and safeguarded *M*_3_were evaluated under each withholding scheme.

### Transfer decomposition

Leave-system-out transfer performance was decomposed into four components:

1. covariate shift,
2. prediction error,
3. query-ranking agreement,
4. calibration behavior. Covariate shift was quantified as ^13^

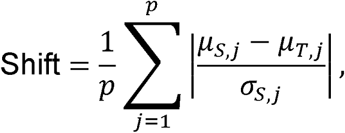

Where *μ_s,j_* and *μ_T,j_* denote source and target means for predictor *j*.

Prediction error was summarized using RMSE,

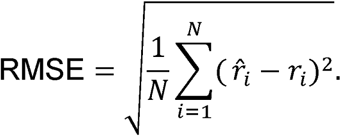

Ranking agreement was measured using Spearman correlation between predicted and observed reliability.

Calibration was summarized using calibration-intercept and calibration-slope errors ^6^.

### Fingerprint similarity and source selection

Fingerprint similarity was evaluated using cosine similarity and correlation-based measures.

Signed-cosine similarity was defined as

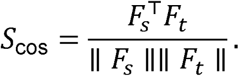

Magnitude-cosine similarity used magnitude fingerprints.

Signed-correlation similarity was calculated as the Pearson correlation between signed fingerprints.

For each target system, candidate sources were ranked by fingerprint similarity and compared with observed transfer performance.

Nearest-source transfer performance was evaluated relative to average-source, context-matched, and random-source baselines.

Leave-one-domain-out analyses repeated the procedure after removing individual fingerprint domains.

### Perturbation-prioritization analyses

For Norman scGPT, perturbations were ranked using reliability estimators transferred from:

1. the fingerprint-nearest source,
2. the fingerprint-most-distant source.

The underlying perturbation predictions remained unchanged.

Observed perturbation performance and enrichment of above-median-performing perturbations were compared among the top-ranked candidates.

### Population-level validation

Population-level prioritization performance was evaluated using 100 constituent-gene-disjoint calibration/evaluation splits for each of ten target systems.

Source-selection strategies included:

- fingerprint-nearest,
- context matched,
- average source,
- random source.

Performance was summarized using:

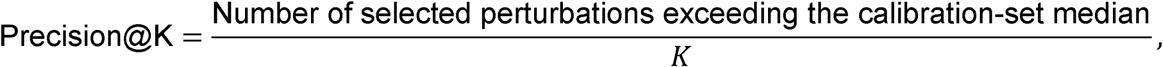

for *K*=10 and *K*=20, and mean observed perturbation performance among the selected perturbations.

These analyses quantified the downstream consequences of reliability transfer for perturbation prioritization.

---

**Supplementary Fig. 1 Pairwise reliability transfer and feature-direction consistency.**

Pairwise source-to-target ROC AUC for binary reliability classification using (A) LASSO, (B) Ridge, (C) Naïve Bayes, (D) Gradient boosting, and (E) Random Forest. Rows denote training systems, columns denote test systems, and diagonal comparisons are omitted. (F) Directional consistency of cellular-component feature associations across model–dataset systems and seven reliability classifiers. Cells show the proportion of evaluable systems sharing the dominant association direction.

**Supplementary Figure 2** A) Binary feature matrix for held-out Norman perturbation queries used in the primary scGPT analysis. Columns indicate query-level binary features and rows indicate perturbation queries. Blue marks feature presence and gray marks absence. Top bars show the number of held-out queries annotated with each feature, and hierarchical clustering groups perturbations with similar feature profiles. (B) Distribution of CollecTRI transcription-factor target burden across the human-gene reference feature set, shown as log10(number of TF targets + 1). (C) Distribution of high-confidence STRING protein-interaction connectivity across the human-gene reference feature set, using STRING v12 interactions with combined confidence score ≥0.9 and shown as log10(degree + 1). (D) Number of human genes annotated with each GO Cellular Component-derived binary feature or ontology-summary feature. Bars show unique genes assigned to each feature category. (E-G) Associations between scGPT prediction performance and continuous query-level features in the held-out Norman test perturbations. Scatter plots show Pearson correlation between predicted and observed delta-expression profiles as a function of (E) log-transformed STRING connectivity, (F) log-transformed CollecTRI TF-target count, and (G) log-transformed total perturbation-level expression. Lines show linear regression fits with 95% confidence intervals; annotations report slope, nominal p-value, and R². **(H)** Associations between curated gene-category membership and prediction performance across model–dataset systems. Rows represent transcription factors, TF targets, tumor suppressors, oncogenes, housekeeping genes and other genes; columns represent model–dataset systems. Red circles indicate higher median performance among category-positive queries, whereas blue circles indicate higher performance among category-negative queries. Circle size reflects statistical strength. Labels report the median performance difference, defined as category-positive minus categorynegative performance, and the corresponding two-sided *P* value.

**Supplementary Figure 3. Population-level prioritization performance across target systems.** Rows show target systems; columns compare fingerprint-nearest, context-matched, average source, and random-source selection. **A,** Precision@20, the fraction of the 20 selected perturbations exceeding the calibration-set median performance. **B,** Mean observed performance among the top 10 perturbations. **C,** Mean observed performance among the top 20. Each cell summarizes the corresponding target-and-strategy result; darker green denotes a larger value. The fingerprint-nearest strategy does not consistently outperform the baselines.

## Declarations

### Ethics approval and consent to participate

Not applicable.

### Consent for publication

Not applicable.

### Availability of data and materials

https://github.com/ads303/scPerturbTrust

## Authors’ contributions

A.S. and T.L. implemented the method, conducted the experiments, interpreted the results. L.Z. inspected the method. S.K. discussed the results and designed the experiments. H.J.P. conceived the project, designed the experiments, and wrote the manuscript.

## Acknowledgements

This research was supported in part by the University of Pittsburgh Center for Research Computing, RRID:SCR_022735, through the resources provided. Specifically, this work used the HTC cluster, which is supported by NIH award number S10OD028483. The authors declare that they have no competing interest.

## EndNote

Import the accompanying RIS file, then use Update Citations and Bibliography. The original numbered reference list is provided in the RIS records.

