## Supplementary figures and images for "Prediction transfer does not imply reliability transfer in single-cell foundation models"

### Supplementary Figure 1

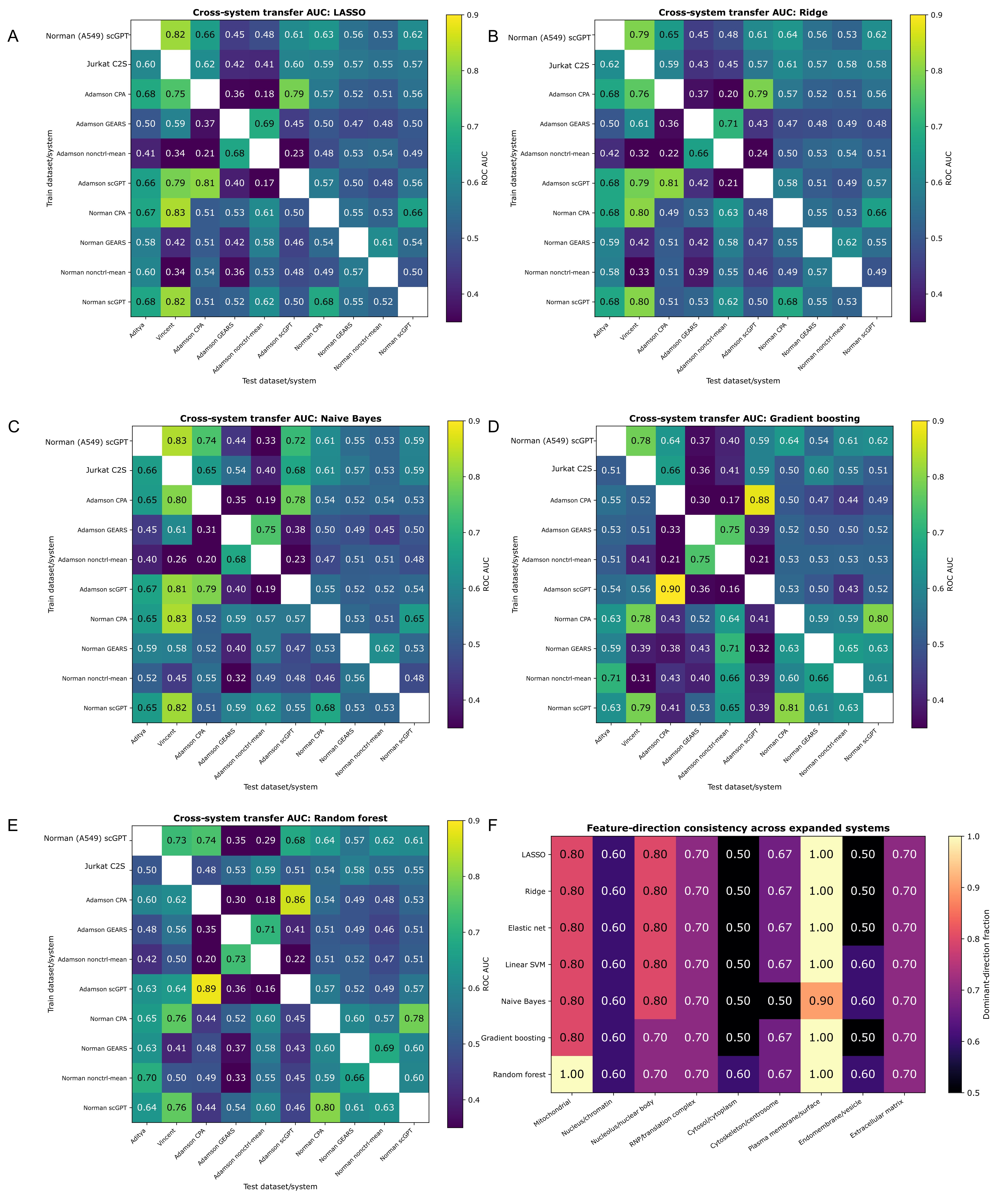

### Supplementary Figure 2

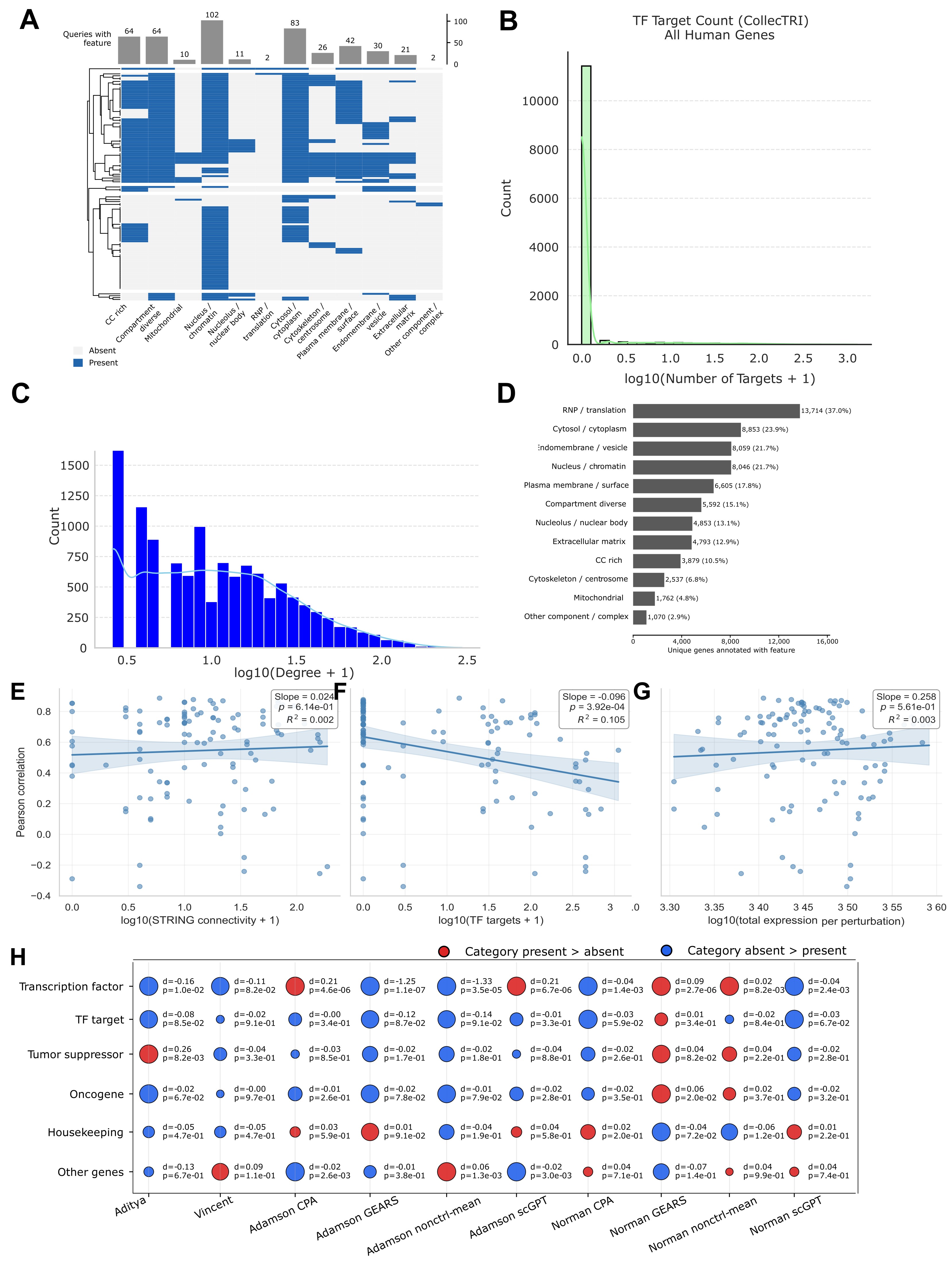

### Supplementary Figure 3

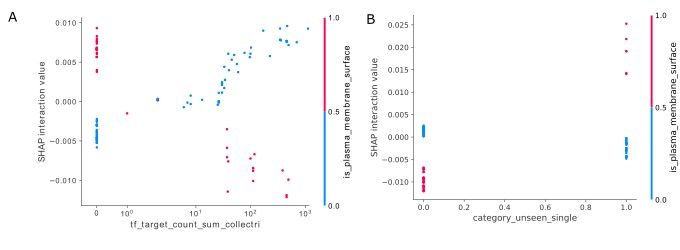
